# OpenMEA: Open-Source Microelectrode Array Platform for Bioelectronic Interfacing

**DOI:** 10.1101/2022.11.11.516234

**Authors:** Gerard O’Leary, Iouri Khramtsov, Rakshith Ramesh, Aidan Perez-Ignacio, Prajay Shah, Homeira Moradi Chameh, Adam Gierlach, Roman Genov, Taufik A. Valiante

## Abstract

Bioelectronic interfaces have the potential to revolutionize the treatment of medical disorders and augment physiology. Implantable devices such as pacemakers and deep brain stimulators have already been deployed to control activity in diseases including Parkinson’s disease and epilepsy. These devices typically operate by delivering electrical stimulation at pre-programmed intervals (known as open-loop stimulation). Recent advances in machine learning and low-power integrated circuits have led to the emergence of personalized medical devices that monitor the user’s state and stimulate in response to measured biological activity (known as closed-loop stimulation). There are two key questions that require fundamental research to achieve breakthroughs in personalized devices: 1) What biomarkers and algorithms are best suited to detecting biological states (e.g. seizures in epilepsy)? and 2) What types of electrical stimuli are optimal for controlling these states? The answer to these questions can be explored *in vitro* using multielectrode array (MEA) systems that interface with biological tissue with reduced experimental complexity, better reproducibility, and fewer confounding variables present in whole organisms. However, existing MEA systems have functional limitations and closed-source designs that prevent researchers from developing improvements. This paper introduces OpenMEA, an open-source platform for closed-loop bioelectronics research. OpenMEA includes designs for the components necessary to build a benchtop *in vitro* laboratory, including electrophysiological recording and stimulation electronics, a microfluidic perfusion system, and physical designs for multielectrode arrays. The system is demonstrated with the electrical recording and stimulation of epileptogenic human and rodent brain slices. The aim of OpenMEA is to democratize bioelectronic research tools to accelerate the deployment of devices for the treatment of disorders and beyond.

## I. Introduction

The use of electromagnetism is ubiquitous in biological systems, with one of the most prominent examples being the action potential in neurons that is mediated by ion channels [1]. Galvani kickstarted the study of bioelectricity when he observed a frog’s leg twitch when struck by an accumulated charge [2], but even after decades of research since, fundamental discoveries are still being made. Elsewhere in nature, it has been uncovered that electrical communication occurs natively in bacterial communities to coordinate metabolic states [3]. Strikingly, emerging research on the “gut-brain axis” is revealing that gut bacteria have a symbiotic relationship with their animal host’s nervous system through bidirectional electrochemical communication to regulate the immune and metabolic systems [4]. It is becoming increasingly evident that bioelectricity is a fundamental mechanism that could be leveraged to treat clinical disorders with further research. *Electroceuticals* have been proposed as the future mainstay of medical treatment. However, it has been emphasized that catalyzing this field requires open innovation and collaborative cross-disciplinary research between neuroscience and engineering in academia and industry [5].

Over the past 25 years, the burden of neurological disorders has increased substantially, with millions worldwide affected by Epilepsy and Parkinson’s disease [6]. Deep brain stimulation (DBS) has been developed as a device-based therapy that can be used to reduce the symptom burden (i.e. seizures in epilepsy and tremor in Parkinson’s disease) where pharmacotherapy is ineffective, or where side-effects are intolerable. DBS uses targeted electrical stimulation to alter underlying pathological brain activity. These devices can be characterized by whether the intervention is triggered at fixed intervals (open-loop), or in response to measured physiological states (closed-loop).

The advent of closed-loop devices has brought with it hope for a radical improvement in the quality of treatment by intelligently responding to pathological brain states. With this approach, stimulation would be reliably delivered precisely when needed and the side effects caused by redundant stimuli could be minimized. However, despite a significant research investment, closed-loop devices have yet to demonstrate a definitive improvement in efficacy over open-loop implants which are still the most commonly used clinically [7]. The utility of these devices is likely predicated on two key unanswered questions: 1) What set of biomarkers and algorithms for pathological state classification are optimal for any given patient? (i.e. *observability*) and 2) What stimulus should be applied to prevent a predicted pathological state? (i.e. *controllability*).

Towards answering these questions, both fundamental research and clinical device validation can be performed *in vivo* and *in vitro*. While *in vivo* validation in animal models is generally required before clinical trials, *in vitro* testing enables the study of biological phenomena with reduced experimental complexity, better reproducibility, and fewer confounding variables present in whole organisms [8]. Cultures of cells can be used, or organotypic slice models of diseases such as epilepsy have been developed to better represent *in vivo* characteristics [9]. The *in vitro* approach has been widely used to assess device design aspects including encapsulation and electrode toxicity [10], closed-loop algorithm development [11], and stimulation protocol evaluation [12].

Microelectrode arrays (MEAs) have played a pivotal role in the *in vitro* study of electrogenic tissues [13]. They are comprised of conductive electrodes patterned on an insulating substrate, and a fluidics chamber (or *well*) which maintains the extracellular media. The electrogenic tissue is either cultured within this well, or thinly sliced brain tissue is added. The MEA is connected to an electronic system which facilitates recording and stimulation of the tissue via the conductive electrodes. This interface can be used to connect and test prototype closed-loop systems with living tissue. However, researchers who wish to integrate proposed advances in hardware (such as custom integrated circuits) or software (such as closed-loop control algorithms) are hindered by insufficient access to the design information of available systems (reviewed in Section II). This is one of the key issues we aim to address with OpenMEA.

Across the scientific and engineering community, there is a growing movement towards *open research*, which supports transparency, sharing, collaboration, decentralization, and accountability [14]. Within this philosophy, o*pen labware* is the free and open sharing of detailed design blueprints for scientific tools and instruments [15] [16]. Examples include open-source 3D printable optics equipment [17] and the Chi.Bio platform for cellular biology [18]. In the context of MEA research, the Potter lab pioneered early closed-loop research with the open-source *Neurorighter* platform [19]. However, in the decade since its development, there have been significant advances in both analog integrated circuits (ICs) and processing capabilities (see Section II). The OpenMEA platform aims to reignite open-source closed-loop electrophysiology with a focus on enabling both fundamental research and implantable medical device development. To put the OpenMEA platform in context, the following section will outline the state of the art in both commercial and academic solutions.

## II. Related Work

The available systems can be categorized into passive and active solutions as outlined in Tables 1 and 2. *Passive* solutions typically only contain conductive electrodes in direct contact with biological tissue. This allows for a wide range of physical properties such as flexibility, transparency, and biocompatibility. *Active* solutions contain nanoscale electronics close to the electrode interface and are typically made from a rigid complementary metal–oxide– semiconductor (CMOS) substrate. These two categories involve a tradeoff between spatial resolution, signal fidelity and experimental practicality (e.g. versatility of materials).

**Table 1.** Commercially available MEA systems.

| Specification | Maxwell MaxOne [28] | 3Brain Bioscan [30] | MCS MEA5000 [32] | Panasonic MED64 [27] | MCS MEA1060 [25] | MCS MEA2100 [26] | **This work: OpenMEA** |
| --- | --- | --- | --- | --- | --- | --- | --- |
| MEA Type | Active | Active | Active | Passive | Passive | Passive | Passive |
| Bottom Perforation | No | No | No | No | Yes | Yes | Yes |
| Closed-loop Processing | PC | FPGA (Intel Arria 10) | PC | PC | PC | Microcontroller (TI TMS320C) | **FPGA (Xilinx Zynq Z-7030)** |
| User Programmable | - | No | - | - | - | Partial | **Yes** |
| Simultaneous Stim/Record | No (Blanking Circuit) | Unknown | Yes (Capacitively coupled) | Unknown | No (Blanking Circuit) | No (Blanking Circuit) | **Yes** |
| MEA Technology | Active CMOS | Active CMOS | Active CMOS | Passive Non-perforated | Passive perforated | Passive Non-perforated | **Passive perforated** |
| Supported Perfusion | Top | Top | Top | Top | **Top and Bottom** | **Top and Bottom** | **Top and Bottom** |
| Electrodes (Rec./Stim) | 26400 | 4112 (4096/16) | 5249 (4225/1024) | 64 | 60 | 256 (max) | 60 |
| Recording Channels | 1024 | 16 | 4225 | 64 | 60 | 252 | 64 |
| Stimulation Channels | 32 | 4 | 1024 | 2 | 2 | 3 | 64 |
| Sample Rate (kHz) | 20 | 20 | 25 | 20 | 25 | 50 | 44.6 |
| Resolution (bits) | 10 | 12 | 14 | 16 | 16 | 24 | 16 |
| Open source | No | No | No | No | No | No | **Yes** |

**Table 2.** Closed-Loop Research Systems.

| Specification | Rolsten et al. 2009 [19] | Müller et al. 2010 [35] | Seu et al. 2018 [33] | Park et al. 2019 [32] | **This work: OpenMEA** |
| --- | --- | --- | --- | --- | --- |
| Processing | PC | FPGA (Xilinx Virtex II Pro) | FPGA (Xilinx Zynq Z-7020) | FPGA (Xilinx Kintex-7) | FPGA (Xilinx Zynq Z-7030) |
| MEA | pMEA (MCS) | CMOS | CMOS | Undisclosed (Up to 128 channels) | pMEA |
| Bottom Perfusion | Yes | No | No | - | **Yes** |
| Electrodes | 60 | 11,011 | 4096 | - | 60 |
| Recording Channels | 64 | 126 | 16 | 2 | 64 |
| Resolution | 16 | 8 | 12 | 16 | **16** |
| Programmable Waveform Gen. | PC-Based | Yes | No | Yes | **Yes** |
| Stimulation Channels | 1 | 42 | 16 (external) | 8 | **64** |
| Sample Rate (kHz) | 25 | 20 | 18 | 32.5 | **44.6** |
| ADC IRN (arms) | 2.11 | 2.4 | 11 | 1.59 | **2.4** |
| Compliance Voltage (V) | +/- 10 | 0-3.5 | +/- 10 | +/- 3.3 | +/- 9 |
| DAC resolution (bits) | 8 | 10 | 16 | 16 | 8 |
| Closed-loop latency (ms) | 63.2 | 0.85 | <2 | - | **<1** |
| Open source | No* | No* | No* | No* | **Yes** |

\*Files not publicly available

Active CMOS arrays can obtain high signal-to-noise ratio recordings by performing signal amplification and digitization close to the electrode before the low-amplitude neural signals are exposed to environmental noise. They can also reduce bulky wiring by means of local time multiplexing, leading to high electrode density which enables sub-cellular resolution. On a practical level, passive MEAs can be based on a broad range of substrate materials (e.g. glass, polymers) which can include mechanical perforations for chemical perfusion. Their fabrication does not require complex CMOS fabrication steps leading to lower customization time and manufacturing cost.

In acute brain slice experiments, recordings are done from the cells at the bottom of the slice. As artificial cerebrospinal fluid (ACSF) is typically drip-fed from a cannula placed over the well, the bottom cells receive less oxygen and nutrients. This causes degrading electrical activity and eventually necrosis. The ability to perform chronic neural interfacing *in vitro* would enable researchers to understand the long-term effects of electrical stimulation and drug interactions [20].

*Passive MEAs* have been developed to mitigate this problem. The inclusion of perforations allows for perfusion to be delivered to the tissue from both top and bottom simultaneously, thereby optimizing the oxygen supply of the slice [21]. Tissue resting on perforated arrays has been demonstrated to stay viable for an average of 10h (or 3.5x) longer than those on non-perforated arrays [22]. Additionally, bottom perfusion creates suction caused by superfusion and capillary wicking [21]. This improves the quality of the electrode-tissue interface and has been shown to increase detectable action potentials by 72% and provide a three-fold improvement in spike detection [18].

*Active MEAs* allow for sub-cellular spatial resolution and have led to significant insights in fundamental neuroscience [23]. There is active research in machining CMOS substrates for applications including perforations [24]. However, due to the computational and experimental complexity of using active arrays, passive arrays have been used more prevalently across neuroscience research.

### A. Commercially available solutions

#### 1) Passive arrays

The most prominent *passive* MEA systems are developed by MultiChannel Systems Gmbh (MCS). In particular, the MEA1060 has been used in a broad range of experiments and had been customized to fit the needs of many research groups [25]. In recent years, the modular MEA2100 was introduced which allows for multiple wells, and significantly include an on-device digital signal processor (Texas Instruments TMS320C6454) for real-time feedback [26]. However, these devices are limited to stimulating only a subset of electrodes using three stimulation channels. Furthermore, the blanking switch system architecture prevents the simultaneous stimulation and recording such as for sub-threshold stimulation. The MED64, originally created by Panasonic, enables recording from 64 electrodes [27]. However, only 2 stimulation channels are supported.

#### 2) Active arrays

The MaxOne (Maxwell Biosystems) offers the highest available density with 26400 electrodes [28][29]. The most significant limitation is the inability to stimulate more than 32 channels, and the system does not support localized closed-loop processing. The BioCAM DupleX (3Brain) offers processing with an integrated Intel Arria 10 FPGA SoC, but users are restricted to the use of pre-designed IP cores (for denoising, filtering etc.) [30]. Furthermore, the *HD-MEA Stimulo*, is limited to 16 channels on a secondary array. More recently, the *HD-MEA Accura* was announced to include 4096 electrodes capable of simultaneous recording and stimulation, but architecture details are not available at the time of writing. Finally, the MEA5000 (MCS) offers the highest number of integrated stimulators but does not include integrated closed-loop processing capabilities.

### B. Published solutions

Several closed-loop MEA systems have been proposed for use with both active and passive arrays. The most significant early closed-loop MEA research was pioneered by the Potter lab with the open-source *Neurorighter* platform [31]. It offered the ability to stimulate all 60 electrodes on a passive array (Multichannel Systems). However, only a single stimulus generator was shared between all electrodes and a PC-based system was used for closed-loop processing which resulted in a best-case latency of >60ms. Recording was facilitated using an external digitizer (NI PCI-6259). In the decade since its development, significant advances in analog IC integration have paved the way for higher fidelity recording, reduced complexity, and reduced cost.

The neural interface system in [32] features 64 input channels (expandable to 128) and 8 stimulation channels with a tightly coupled FPGA for signal processing. The 64 input channels use two stages of input switches with a single time multiplexed 16-bit SAR ADC (TI ADS8422). It should be noted that although the ADC offers an input referred noise of 1.59μV, the addition of the two multiplexing stages is not reported. Furthermore, the stimulation capabilities are limited to 8 channels using 16-bit resolution DAC. Closed-loop processing is supported using a Xilinx Kintex-7 FPGA. Two significant CMOS MEA-based closed-loop systems have been reported (Table 2), although these are not suitable for chronic applications. The system presented in [33] adopts 3Brain’s 4096-electrode CMOS technology [34]. The approach offers high-density recording with high-performance processing but relies on an external stimulus generator (*Plexon PlexStim*) which is limited to 16 electrical stimulation channels and does not support real-time programmable waveforms. Another collaborative approach is outlined in [35], where a CMOS MEA developed by Maxwell Biosystems [36] was combined with an FPGA based system to provide closed-loop control. However, the Xilinx Virtex II FPGA provides limited scope for further expansion. The open-source transparent MEA described in leverages the OpenEphys platform, but it does not support stimulation or closed-loop processing. Note that no existing solution fulfills all of the following requirements:

1. Open-source design to allow the integration of custom hardware and software.
2. Supports perforated MEAs for chronic experimentation.
3. Includes an FPGA for prototyping synthesizable logic for ASIC-based neural interfaces.
4. Supports an embedded operating system to ease the integration of new peripherals.
5. Supports simultaneous recording and stimulation.
6. Supports the real-time adaptation of neuromodulation waveforms in response to inferred states.
7. Allow stimulation to continuously be applied to a subset, or all electrodes simultaneously.

Based on these shortcomings, the development of the OpenMEA platform will be outlined in the following section.

## III. Overview

OpenMEA (Figure 1) aims to condense the features of a typical lab MEA setup into a benchtop form factor. In doing so, it should also facilitate the use of additional instruments such as a fluorescence microscope and should protect the system from external environmental noise. As will be detailed in the following sections, the platform includes the key necessary components for *in vitro* bioelectronics research including: 1) passive microelectrode arrays and tissue chambers 2) electrical recording and stimulation hardware 3) a perfusion system for extracellular fluid and pharmaceutical delivery and 4) processing capabilities for real-time analysis and closed-loop stimulation.

**Figure 1.**
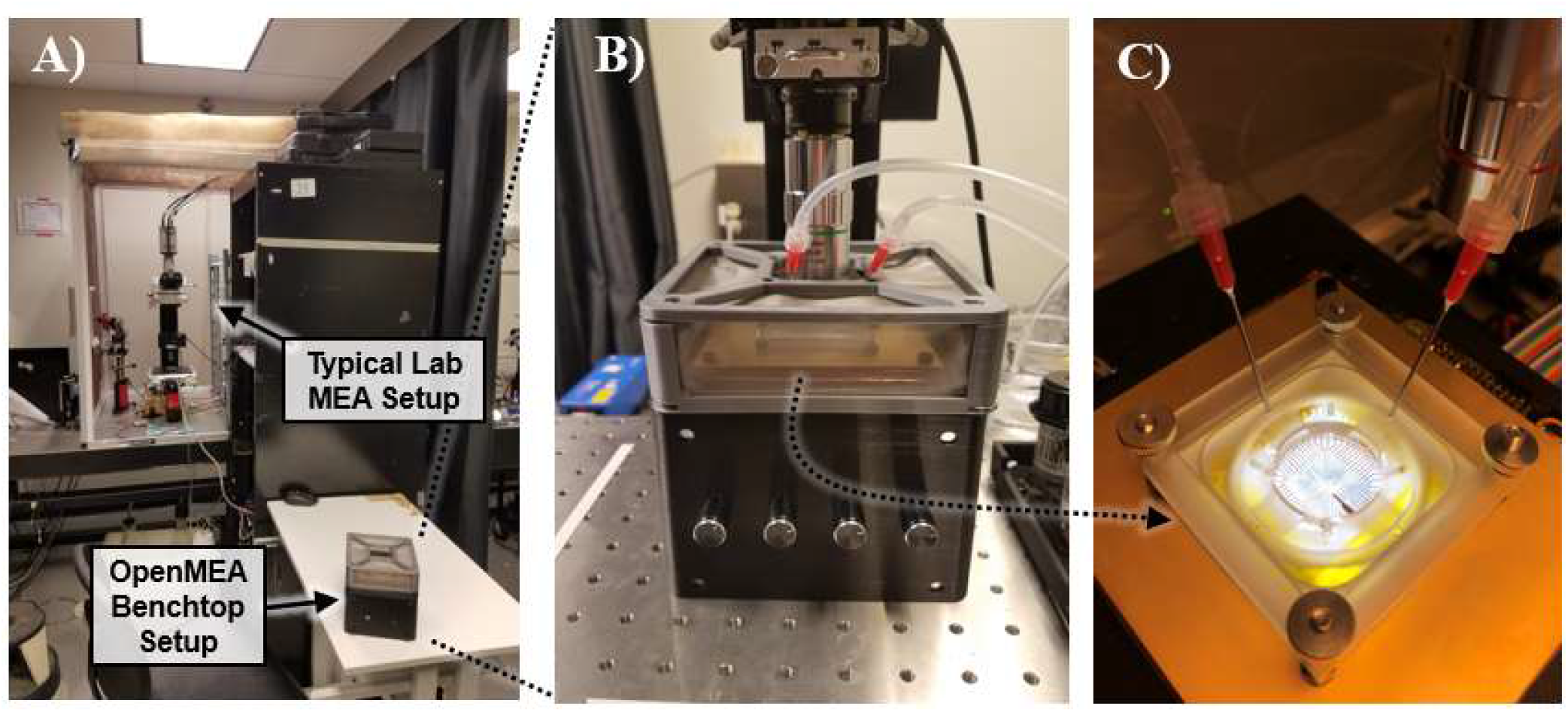
OpenMEA Overview. A) OpenMEA is a benchtop system in contrast to a typical large laboratory setup. B) System configuration with fluorescence microscope for calcium imaging. C) MEA with an integrated perfusion system for the delivery of artificial cerebrospinal fluid (ACSF) and pharmaceuticals.

In comparison to the state of the art in Table 1 and Table 2, OpenMEA offers the highest performance user-programmable processing capabilities with the Xilinx Zynq Z-7030. It is one of the only systems that supports perforated MEAs with top and bottom perfusion for long-term experimentation. Its recording capabilities are on par with many commercial systems with 16-bit 44.6 KHz sampling on up to 64 channels. Furthermore, each electrode has a dedicated stimulator with 8-bit resolution, allowing for independent control with unrestricted waveform morphologies. This contrasts with other systems which are limited to bi-phasic waveforms on subsets of electrodes.

During a typical experiment, the electromagnetic shield is removed, and biological tissue is placed on the MEA. The top and bottom perfusion and suction are connected, and the media flow rate is calibrated. The shield is then replaced, and the processing system initializes the headstage electronics. Activity within the tissue can then be recorded, and stimulation can be enabled using a graphical user interface.

Due to the broad range of possible experiments and models that should be facilitated (from electroactive cells to organotypic slices), the ability to modify the platform is paramount. The design files have been released online (OpenMEA.github.io) under a *Creative Commons Attribution-ShareAlike 4*.*0 International License* [38]. This allows users to copy, modify and redistribute the design for any purpose, including for commercial use. However, users must redistribute their iterations under the same license, give appropriate credit to the source, and may not apply legal measures such as patents to restrict others from following the same license. These restrictions do not apply to the software and device prototypes that use the OpenMEA platform for validation. For example, medical device developers may integrate prototype hardware with OpenMEA to perform *in vitro* testing without concerns about the final product being subject to licensing restrictions.

## IV. Microfluidic media perfusion system

Experiments with biological tissue require the use of a fluidic chamber to maintain the extracellular media. In the case of acute brain slices, artificial cerebrospinal fluid (ACSF) is drip-fed from a cannula placed over the well during electrical interfacing (Figure 2A). As we will discuss, the perfusion delivery method has implications for multi-modal neural interfacing.

**Figure 2.**
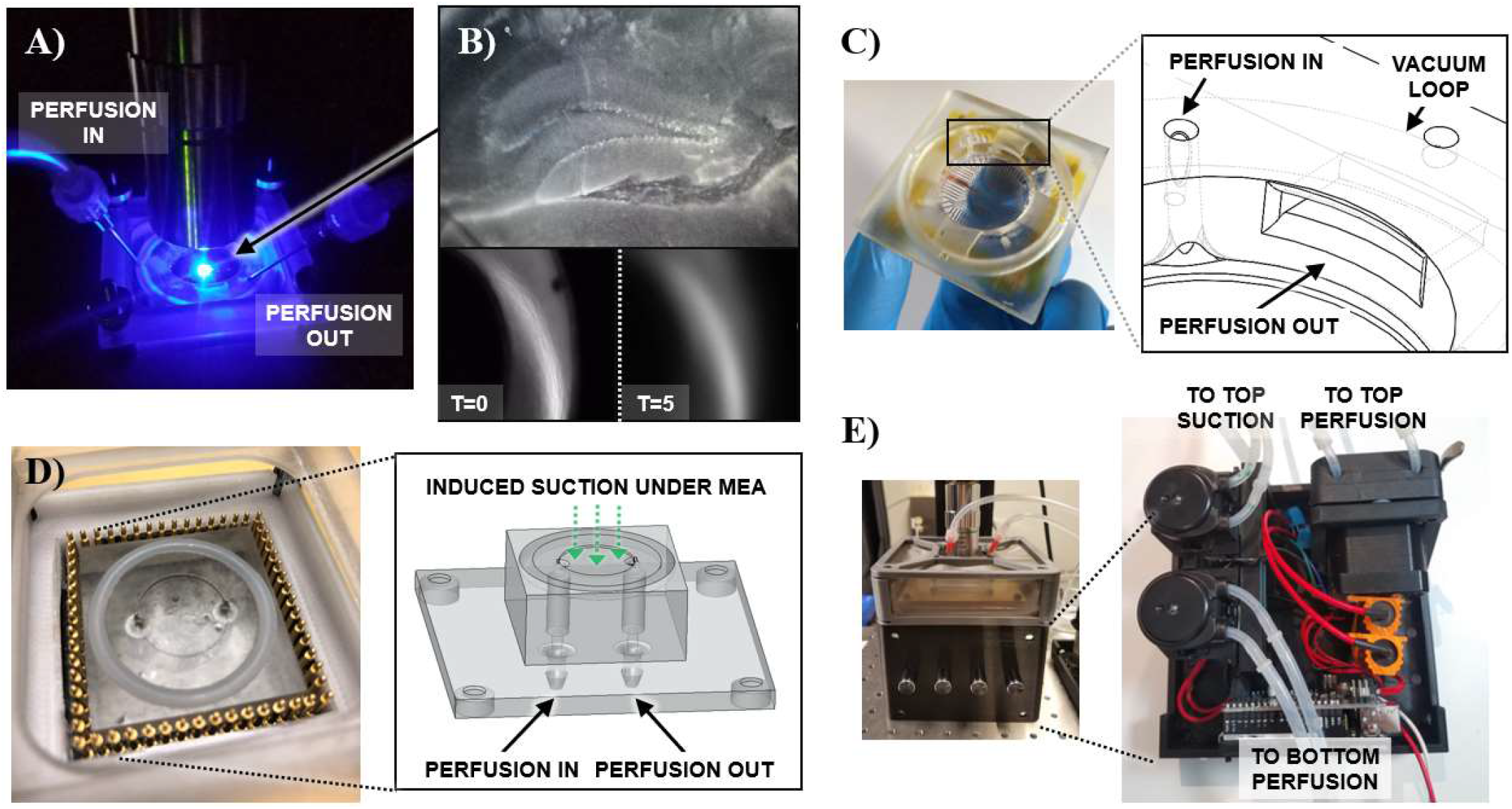
Imaging-optimized perfusion microfluidic methodology. A) Overview of perfusion setup during calcium fluorescence imaging. B) Rodent brain slice image and an associated issue with typical top perfusion approaches. At T=0, the fluorescence optics are calibrated. At T=5, the image has lost focus. C) To circumvent focusing issues, an integrated microfluidics channel delivers incoming media to the bottom of the well, and an overflow opening channels outgoing media to a vacuum loop. D) Bottom perfusion microfluidics bracket containing a cavity with input and output ports under the MEA perforations. Spring-loaded pogo-pins provide electrical contact between the MEA contacts and neural interface electronics. E) Perfusion system consisting of programmable peristaltic pumps for precise delivery of top perfusion. The front panel includes potentiometers for manual calibration of flow rates.

There is a growing trend towards the simultaneous use of optical imaging and stimulation with electrical interfacing. For example, this combination enables the exploration of electrical stimulation parameters including amplitude and frequency while visually assessing neural activation dynamics [39]. During such experiments, it is essential to maintain a constant focus and avoid imaging distortion which can hinder the interpretation of data. A calcium imaging experimental example is illustrated in Figure 2B with a transgenic rodent coronal brain slice of the dentate gyrus (Jackson Laboratory, Thy1-GCaMP6s). Nonidealities in the delivery of top perfusion causes fluctuations in the ACSF level, leading to focusing issues. Using a nearby OpenMEA electrode (Section V) as a reference point, at T=0, the fluorescence optics are calibrated. At T=5, the image has lost focus due to an unstable perfusion level caused by the inhibition of suction by ACSF surface tension.

To circumvent this, the OpenMEA well includes design features to regulate the flow of media (Figure 2C). This is achieved using both a microfluidic channel to deliver media to the bottom of the chamber, and a vacuum loop around the MEA perimeter. The vacuum loop extracts depleted ACSF from the central chamber via an overflow slot using gravity as opposed to direct suction. This combination minimizes both optical focusing issues and droplet-induced motion artifacts in electrical recordings. The inlets to the input channel and vacuum loop are connected via a cannula and silicone tubing to an external perfusion system.

As outlined in Section II, perforated arrays with bottom perfusion prevent necrosis in cells recorded at the bottom of slice and enable suction to increase the physical coupling between tissue and electrodes. In OpenMEA, the perforated arrays described in Section V are placed on a bottom bracket which enables the suction of depleted media through the array, and the superfusion of fresh media into the well (Figure 2D). This bracket contains a microfluidic cavity with input and output ports under the MEA perforations. The bracket is spring-pressed to a silicone O-ring to prevent leaks between the perfusion assembly and the active electronics.

Both the top well and bottom bracket are manufactured using a masked stereolithography apparatus (MSLA) 3D printing process (Elegoo Mars 2 Pro) with a transparent UV photosensitive polyurethane acrylate (PUA) resin (Siraya Tech, Clear). PUA has been demonstrated to have low biotoxicity (enabling *in vitro* use), and low conductivity (preventing unintended electrical coupling with electrode contacts) [40]. While microfluidic-based perfusion has recently been demonstrated in the domain of cell-culture research using advanced fabrication methods [41], OpenMEA leverages a more recently accessible additive manufacturing process with increased design capabilities. Furthermore, this approach allows for eased design customization for applications such as multi-well experimentation, and reproducible designs among research groups.

OpenMEA also includes an integrated multi-channel perfusion pump system with programmable and manual flow rates (Figure 2E). The system uses peristaltic pumps in which a motor-driven rotor with rollers attached to its circumference are in compressive contact with flexible silicone tubing. As the motor shaft rotates, the rollers compress the flexible tube, positively displacing the tube contents as they rotate by. This process, known as peristalsis, results in positive pressure at the pump output, and negative pressure at the pump input. The type of motor used to drive the rotor influences three parameters: the maximum pressure, the controllability, and the smoothness of delivery. DC motors achieve smooth rotation with a high rate of rotation (and hence pressure). Stepper motors enable precise control of fluid movement which is useful for setting specific ACSF flow rates and titrating the delivery of drugs.

An ACSF flow rate of 2-3ml/min is recommended for top perfusion to maintain brain slices during experiments [42]. OpenMEA uses a stepper motor-based pump to ensure precise control over a flow range of 0-140ml/min for top perfusion delivery (Fafeicy, B08B1MNTFD). For top suction and bottom perfusion, a DC motor-based pump is used to provide a high flow rate and smooth suction (Gikfun, AE1207). The motors are driven using H-Bridges (Toshiba TB6612) which are switched to control speeds using a PWM IC (NXP PCA9685) and an ATmega328P microcontroller. The bill of materials is <$200 in contrast to commercial alternatives which are an order of magnitude higher.

## V. Microelectrode Arrays

The most broadly used MEAs for *in vitro* neuroscience experiments are typically manufactured using biocompatible conductive materials (e.g. indium tin oxide, titanium nitride) deposited on a polyimide-on-glass base [43]. Attractive property of these arrays includes transparency (permitting easy observation of tissue), low toxicity and electrode sizes of <100µm (enabling spatially precise interfacing). However, as outlined in Section II, the rigid materials used hinder the ability to machine perforations for bottom perfusion. Furthermore, the expense of materials and manufacturing results in a cost to the end user of >$500 per array and must be replaced typically after 30 uses. A key question then arises: do more optimal and accessible manufacturing alternatives exist while providing small enough feature sizes for high spatial resolution?

### A. Optimal electrode sizing

The level of required spatial resolution is highly application dependent. For experiments involving single-neuron interfacing, sub-cellular resolution offers the ability to study the propagation of action potentials [23]. For intuition on an appropriate electrode size, pyramidal cells are the most common neurons in the cortex and have cell bodies that range from 20–120 µm in diameter [44]. Studies have found the optimal electrode diameter for the measurement of extracellular action potentials (EAPs) to be between 30-50μm, but note that noise and signal attenuation depend more on the electrode impedance than on electrode size [45]. EAPs from individual neurons in the frequency range from 300 Hz to 5 kHz, with amplitudes of 10s of microvolts [46]. Electrophysiology is also performed at larger scales, where ionic currents caused by the activity of populations of cells can be measured in the form of local field potentials (LFPs) [47]. These signals are in the frequency range from 1 Hz to 300 Hz with amplitudes on the order of microvolts to a couple of millivolts [46]. LFPs have become increasingly popular over the last decade as they offer multiple insights into the mechanisms underlying cortical processing at a network level [47] and have been used as biomarkers for adaptive deep brain stimulation (DBS) in Parkinson’s disease [48] and in epilepsy [49]. In the case of DBS, large cylindrical macroelectrodes with surface areas of ∼6 mm^2^ are used to maximize the activated volume of brain tissue for therapeutic stimulation. So what is the appropriate macroelectrode size for *in vitro* MEA experiments?

For intuition, the rodent brain typically used in acute in vitro neuroscience experiments is approximately 35x smaller than human, with lower neuronal density [50]. An estimate on the scaled electrode area would therefore be 0.17 mm^2^, or a circular geometry with a diameter of ∼500µm. The use of large geometries not only recapitulates the clinical DBS context, but also eases design requirements for both manufacturing and electronic circuits, as larger surface areas have a lower impedance. Furthermore, single-cell activity can be concurrently analyzed using optical imaging, and spatially-precise stimulation can be achieved using emerging methods such a virtual electrodes [51].

In summary, single-unit interfacing and population-level interfacing present distinct and complementary opportunities to interact with electrogenic tissue. However, each application has an optimal electrode configuration (approximately 50µm and 500µm, respectively). OpenMEA therefore includes both *micro-* and *macro-electrode* 60-channel arrays optimized for single-unit interfacing and LFPs (Figure 7).

### B. OpenMEA multielectrode arrays

The use of printed circuit boards (PCBs) for lab-on-chip applications was first suggested many years ago but has recently re-emerged due to the increase of accessibility and affordability of advanced manufacturing capabilities [52]. Of relevance to *in vitro* interfacing, such capabilities include the ability to produce high density features (<100µm) with biocompatible materials such as electroless nickel immersion gold (ENi/IAu). Transparent, flexible polymer substrates such as polyethylene terephthalate (PET) can be used, which have demonstrated low toxicity *in vitro* [52]. Furthermore, multi-layer PCB technology enables 3-dimensional features, such as surrounding recording electrodes with a large reference electrode for increased recording quality. OpenMEA leverages these capabilities to produce functional MEAs at a cost of <$1 per array.

The EAP MEA (Figure 3A) uses a feature size of 50µm with a 750µm pitch, and the LFP MEA (Figure 3B) uses a 500µm perforated via electrode with a 1250µm pitch. The board stackup consists of a PET core with double-sided copper and a transparent liquid photo imageable (LPI) polymer coverlay. An ENi/IAu plating process (38µm Ni, 0.2µm Au) is used to prevent oxidization and improve biocompatibility. For the final MEA assembly, the PCB top-layer contacts are covered with polyimide tape to minimize electrode coupling after being adhered to the perfusion well (Section IV) using FDA grade silicone (Silco, Sil-Bond RTV 4500). Spring-loaded pogo-pins provide electrical contact between the MEA contacts and neural interface electronics (Figure 2D).

**Figure 3.**
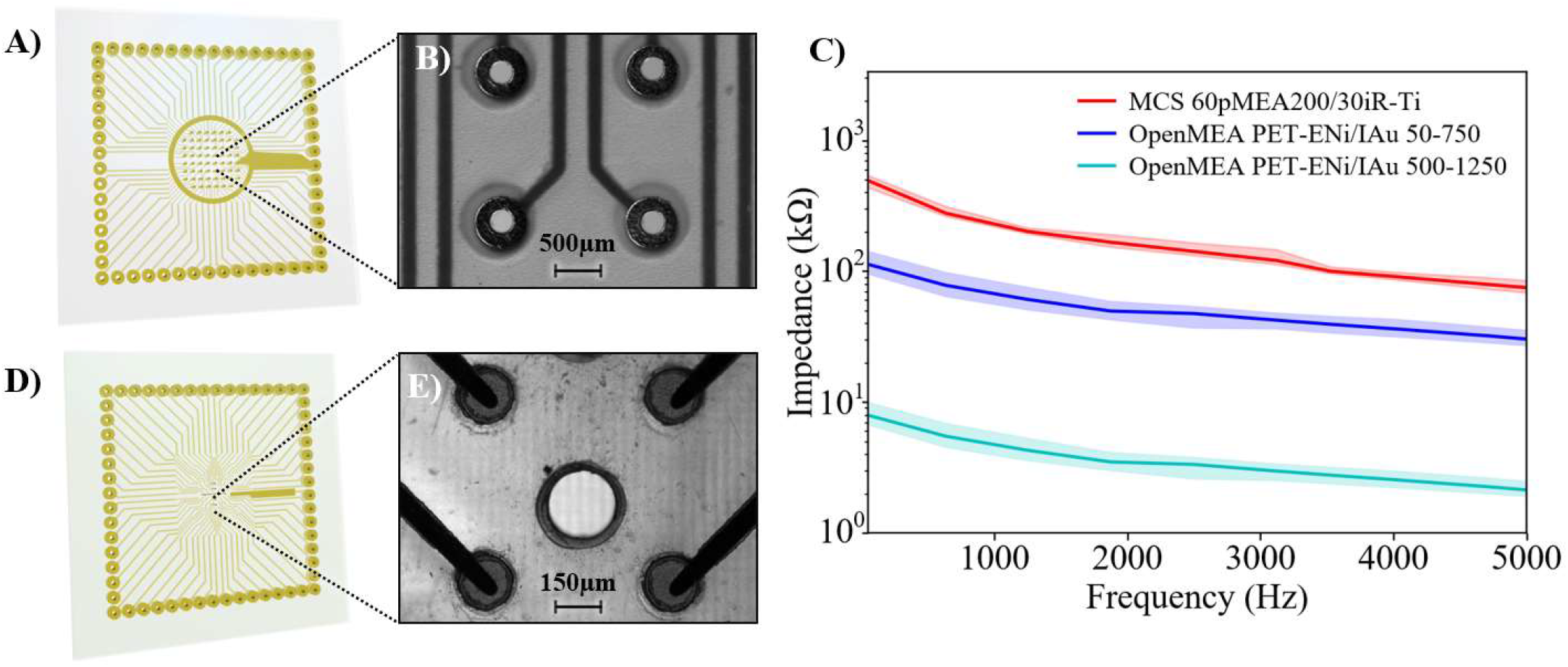
OpenMEA microelectrode arrays. A) Transparent PET-ENi/IAu printed circuit board-based (PCB) 60-macroelectrode array includes media perforations. B) Microscope image of array at 20x magnification. C) The impedance of the OpenMEA electrodes are 5kΩ and 79 kΩ at 1KHz compared to 275kΩ with commercially available arrays. D) Transparent PET-ENi/IAu MEA. E) Microelectrode array at 20x magnification.

As illustrated in Figure 3C, the electrode impedance of the LFP PET-ENi/IAu OpenMEA is ∼5kΩ at 1KHz compared to 275kΩ with the most widely used commercially available MEA (Multichannel Systems MCS 60pMEA200/30iR-Ti). This is due to the increased surface area of the electrodes both because of a larger diameter, but also due to the use of plated holes which effectively serve as 3D electrodes. Impedance measurements were performed in a saline solution using a 50-5000Hz frequency sweep (Intan RHS Stim/Recording Controller).

## VI. Modular electronic recording and stimulation

The OpenMEA electronics for neural signal acquisition and stimulation are implemented on segregated printed circuit boards. Noise sensitive electronics are located inside the shielded headstage (Section VII) close to the source, and digitized samples are transmitted using low-voltage differential signaling (LVDS) to a distal processing system. In this case, noise is minimized as the device leads carry digital data rather than noise-susceptible analog signals.

The OpenMEA headstage mainboard (Figure 4A) contains spring-loaded pogo-pins for electrical contact with the MEA. the active neural interface electronics modules and an LVDS interface header. The analog interface module (Figure 4B) is responsible for the amplification and digitization of neural activity, along with stimulation to induce neural activity. The module is designed around the Intan RHS2116 neural interface IC which contains an array of 16 stimulation/amplifier blocks. Each channel includes a low-noise amplifier with programmable bandwidth and a constant-current stimulator with an 8-bit programmable amplitude to support arbitrary waveforms. The circuit architecture combines stimulators, amplifiers, analog and digital filters, a multiplexed 16-bit analog-to-digital converter (ADC), and a flexible electrode impedance measurement module on a single silicon chip. A low-distortion, high-speed analog multiplexer (MUX) allows all the amplifiers to share one on-chip ADC. The ADC can sample each channel up to 44.6 kSamples/s. A standard module form factor allows for interchangeability with a user’s custom analog interface ICs.

**Figure 4.**
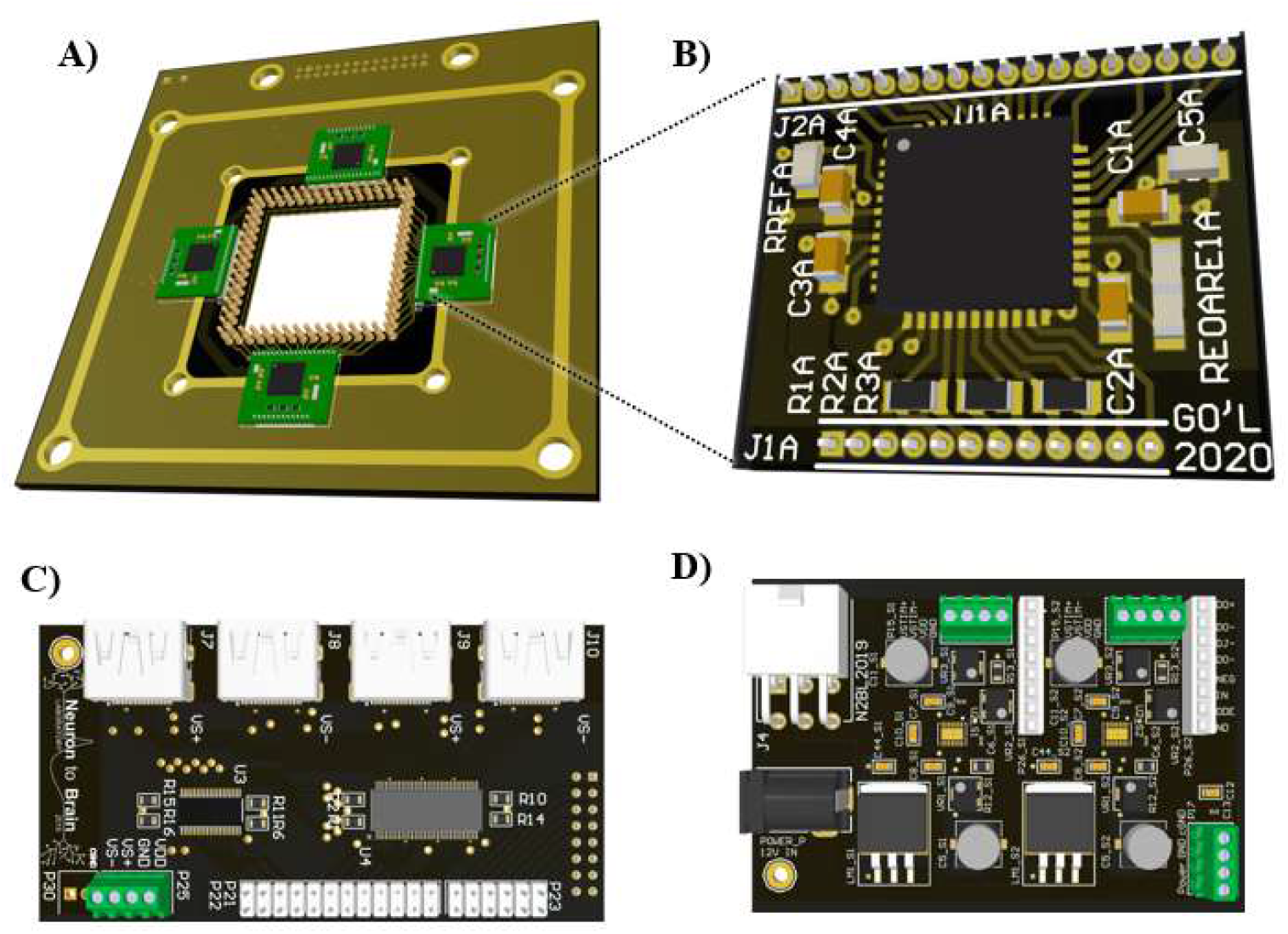
OpenMEA modular recording and stimulation electronics. A) Headstage mainboard containing the active neural interface electronics modules and spring-loaded pogo-pins for electrical contact with the MEA. B) Neural interface IC module for 16 channel recording and stimulation. C) Low-voltage differential signaling (LVDS) interface board for communication between the headstage and the processing system. D) Power supply board for system and stimulation voltage generation.

The OpenMEA design also includes support boards for power and data transmission. Digital communication between the headstage and processing system (Section VIII) is enabled using a serial peripheral interface (SPI) over LVDS interface. The interface board (Figure 4C) hosts noise-tolerant LVDS transmitters (TI SN75LVDS387) and receivers (TI SN65LVDT388). The power supply board contains the circuits responsible for system voltage and stimulation voltage generation (Figure 4D). A low-dropout linear voltage regulator (LDO) is used for minimal noise at the sample acquisition IC (TI LM1085), and high-voltage positive and negative stimulator supplies are generated using a dual-polarity LDO with an inverting charge pump (ADI LTC3260).

## VII. Dual-layer electromagnetic shielding

One of the greatest challenges in recording from electrogenic tissue is obtaining a strong signal-to-noise ratio (SNR). In the case of neural tissue, signals amplitudes are in the range of 10s of microvolts for action potentials, and millivolts for local field potentials [46]. Large sources of electromagnetic noise can therefore hinder recordings. The most significant environmental electrical noise is related to the AC delivery of mains power with a fundamental frequency of 50/60Hz and additional harmonics [53]. Digital processing devices and embedded systems generated noise due to due to internal switching, which can be radiated as electromagnetic emissions, or coupled through a conductor to recording systems [54]

To avoid ambient noise, a controlled electromagnetic environment can be used in the acquisition of bioelectric signals. A Faraday cage is an electromagnetic enclosure with a zero internal electromagnetic field [55]. It achieves this by redistributing the electric charge of the external electric field within the enclosure’s conducting material, which in turn cancels the field’s effect in the cage’s interior. The OpenMEA design uses a dual-layer Faraday cage approach to minimize noise interference (Figure 5). Layer 1 consists of an FDM 3D printed outer cage with stainless steel mesh. The internal layer is a PCB-based solid 4 oz copper shield which encapsulates the analog sensing electronics. This combination is demonstrated to reduce 60 Hz and harmonic noise power by x16 (−24 dB).

**Figure 5.**
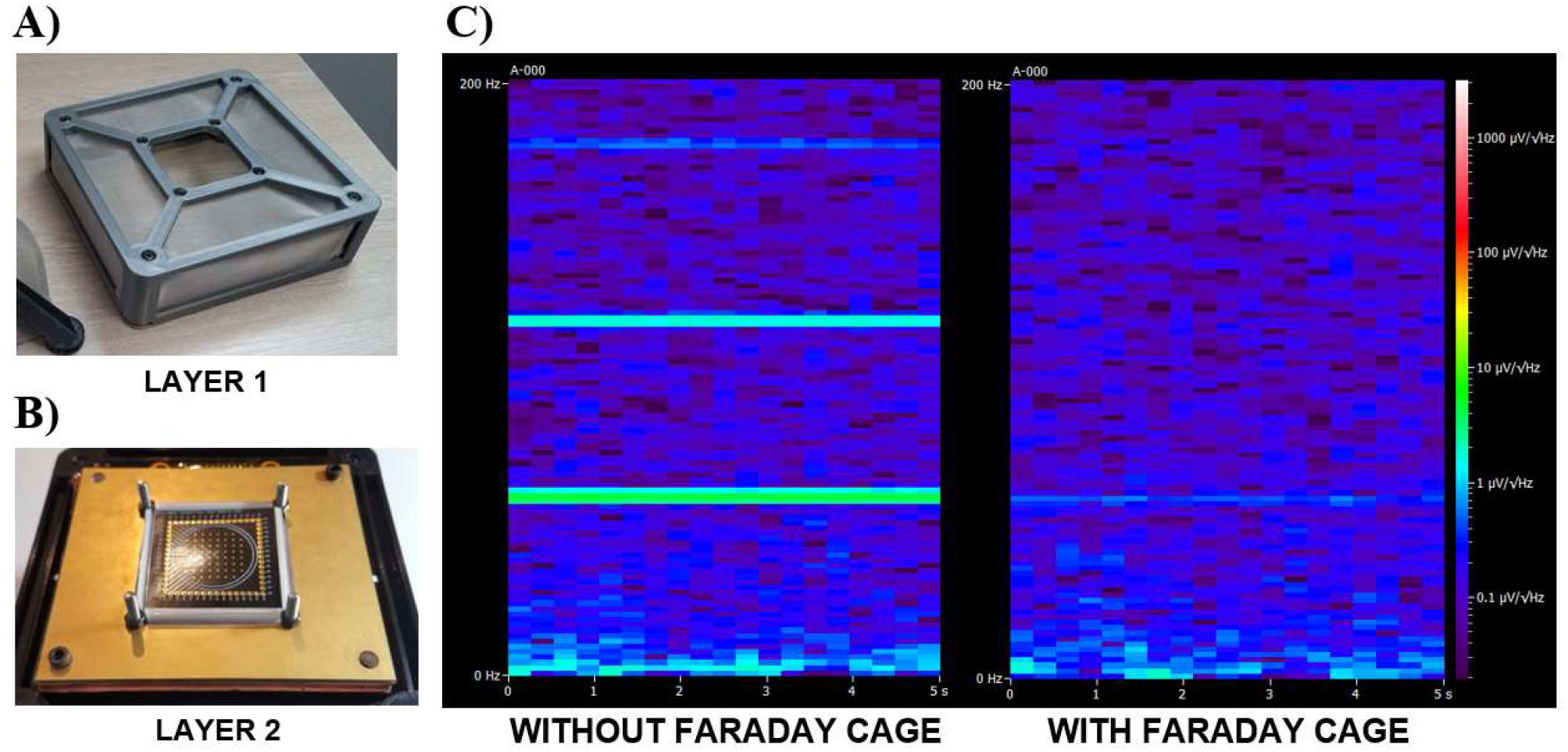
Integrated dual-layer Faraday cage for electrical noise shielding. A) FDM 3D printed outer cage with stainless steel mesh. B) Internal PCB-based solid 4 oz copper shield which encapsulates analog sensing electronics. C) The dual-layer Faraday cage reduces 60 Hz and harmonic noise power by x16 (−24 dB).

In a previously published design iteration [51] noisy electronics were implemented on the same PCB as sensitive analog ICs. The noise aggressors in the system would ideally be placed away from the recording electrode, but due to the low amplitude of the signals acquired by the MEA, the AFE must be placed as close as possible to the electrodes. For this reason, OpenMEA uses segregated PCBs, where sensitive electronics are located inside the shielded headstage as close as possible to the source, and digitized samples are transmitted using low-voltage differential signaling to a distal processing system (Section VIII).

## VIII. Data processing and computation system

Data processing and computation are critical components in closed-loop neuromodulation research. This includes supporting the development of state detection and adaptive stimulation algorithms [11], along with automating experimental protocols towards “self-driving laboratories” [56]. OpenMEA provides two tiers of programmability: software-based interfacing using *OpenMEA Studio* and low-latency hardware-based interfacing using a system-on-chip field-programmable-gate array (SoC FPGA). Alternatively, for basic recording and stimulation functionality, the system is compatible with commercially available controllers (e.g. Intan RHS Stim/Record System).

### A. Hardware-based processing system

The hardware-based processing system is designed to address three requirements that arise in closed-loop neuromodulation research: 1) high-throughput, 2) low-latency and 3) device translatability.

A major bottleneck in scaling the number of electrodes in neural interfaces is the processing *throughput* requirements for the increasing number of digitized sample streams. This generally involves signal filtering, spike detection, and identifying the neuron from which a detected spike was generated (spike sorting). This activity can then be used to infer a network state, which in turn is used to select a stimulus in response (i.e. closed-loop stimulation). This inference stage is application dependent, and so flexibility is needed to allow researchers to define both a state of interest from detected spiking activity, and the desired spiking activity to be induced.

The required processing *latency* in closed-loop experimentation is experiment dependent. For intuition, action potentials have a typical duration of about 1ms and event related potentials (ERPs) typically have timescales on the order 100ms [57]. For experiments involving spike-timing-dependent-plasticity (STDP), processing latencies on the order of <10ms are necessary to ensure that state-responsive stimuli are applied before becoming outdated [58]. In general, low latency processing enables tighter coupling between measuring and responsively perturbing electrogenic tissue.

The ability to develop closed loop processing algorithms that can be *translated* from software on a general-purpose computer/server to a power-constrained implantable medical device is highly desirable. As will be described in Section X, implantable devices use application-specific integrated circuits (ASICs) to meet the constraints associated with long-term battery operation. Digital processing on such ASICs is typically implemented using a hardware description language (HDL).

OpenMEA uses an SoC-FPGA (Xilinx Zynq Z-7030) platform which enables the use of HDLs (e.g. Verilog) to prototype designs in programmable logic (PL) and ease the translation of algorithms from the benchtop to an implant. A general-purpose dual-core processing subsystem (PS) sits alongside the FPGA fabric, enabling embedded Linux support (PetaLinux). This opens a range of closed-loop processing capabilities including the use of existing peripheral and machine learning development frameworks.

The firmware on OpenMEA’s System on Chip (SoC) serves as the bridge between the OpenMEA Studio and hardware. The PL contains a controller module to interface with the neural interface headstage over SPI, memory-mapped registers for configuring the controller, and dual-channel FIFOs for each neural ADC/DAC (one channel for the incoming neural data, and another for outgoing stimulation commands). The PS runs higher-level processing and abstracts the low-level networking commands necessary to transmit neural data to send samples and receive stimulation commands from the host computer. Ethernet communication applications run in software to packetize streams of neural data from the headstage control FIFOs, and to de-packetize streams of stimulation commands from *OpenMEA Studio* using the user datagram protocol (UDP). Custom Linux kernel modules are used to interface with logic present on the PL side of the SoC.

### B. Software-based processing system

The user interface for OpenMEA is a desktop application available for Windows, Mac, or Linux (Figure 7). The application allows controlling the OpenMEA device, viewing and saving real-time electrophysiology data from OpenMEA, and stimulating any number of electrodes. The application is made to be easily extensible, with the ability to add custom closed-loop stimulation processing.

When collecting the electrophysiology data, the user can change the sampling rate, apply software bandpass or 50/60 Hz comb filters, and view the spectrogram for any channel. The user can also save the live data in Neurodata Without Borders (NWB) format for further off-line analysis [59].

The application also allows the user to stimulate any number of electrodes with arbitrary waveforms defined in Wave Audio File Format (.wav files). These files can be of arbitrary length and the application allows to loop the files indefinitely. Any number of electrodes can be stimulated concurrently with different waveforms. For convenience, the application also provides a user interface to generate short biphasic pulses.

The application’s UI is written using TypeScript and Electron, and the back end is written in Python. The structure of the Python back-end allows the users to easily extend or modify the application after it has been installed, including adding custom neural signal processing, closed-loop stimulation algorithms. As demonstrated in Figure 6, such processing can be implemented using an average latency of <100ms.

**Figure 6.**
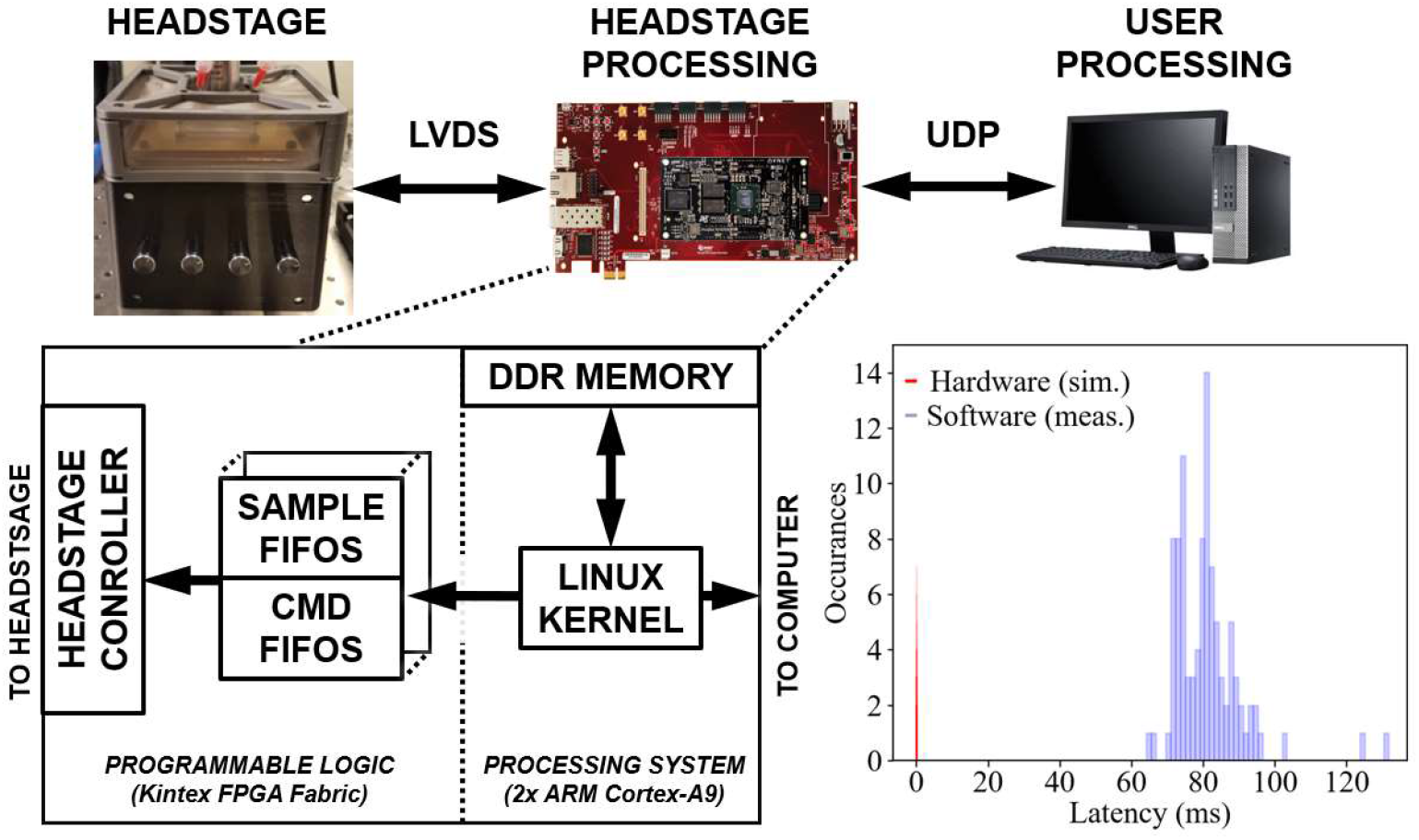
Processing system. A) Data flow diagram from headstage to user interface. B) FPGA block diagram for Zynq-based processing system. C) Histogram for closed-loop processing showing an 80ms latency for software-based experiments and 100ns for hardware-based implementations.

**Figure 7.**
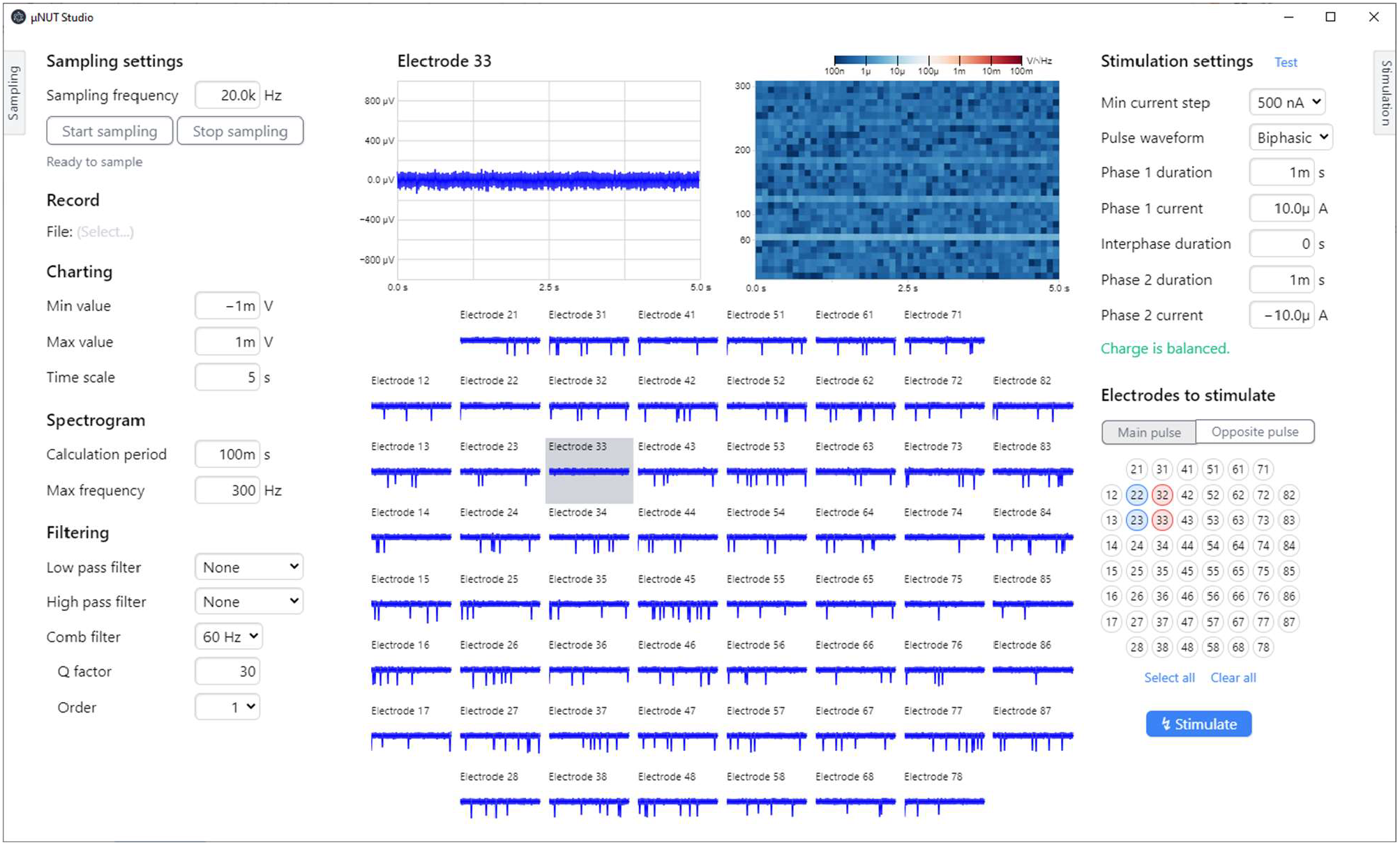
OpenMEA Studio allows for viewing, analyzing, and saving neural recordings, controlling stimulation, and implementing custom closed-loop processing functionality.

## IX. In vitro validation

The OpenMEA system was validated using an *in vitro* experimental setup with both human and rodent brain tissue. Human cortical tissue was acquired during an anterior temporal lobectomy on patients undergoing resective surgery for treatment of pharmacoresistant epilepsy with informed consent at the Toronto Western Hospital. A 1 cm^3^ block resected from the middle temporal gyrus (MTG) was sectioned into 500µm thick slices using a vibratome (Leica VT1200S). Rodent brain slices (Jackson Laboratory, Thy1-GCaMP6s) were prepared as outlined in [60] to produce 500um thick coronal slices from the somatosensory cortex. Briefly, after injecting mice with 0.1% euthanyl, a heart perfusion was performed before decapitation and the brain was rapidly extracted and cut in ice-cold oxygenated sucrose solution. Slices were then left to recover in standard ACSF at room temperature for 60 minutes and incubated.

Prepared slices were transferred to the OpenMEA headstage for interfacing where it was perfused at ∼3mL/min (top perfusion) with carbogenated ACSF (Figure 8A). Epileptiform activity was induced using 4-aminopyridine (4-AP; 50 µM), a non-selective potassium channel blocker which extends the duration of action potentials by preventing cell repolarization, and causes increased synchronous network activity [61]. As illustrated in Figure 8B, ictal bursting activity can be seen originating in electrode 52 with apparent spatial propagation to the adjacent electrode. This is in line with experiments conducted in literature [62]. Figure 8C illustrates the interruption of ictal discharge activity using manually initiated electrical stimulation (10 pulses with a frequency of 100 Hz and a 100µA amplitude).

**Figure 8.**
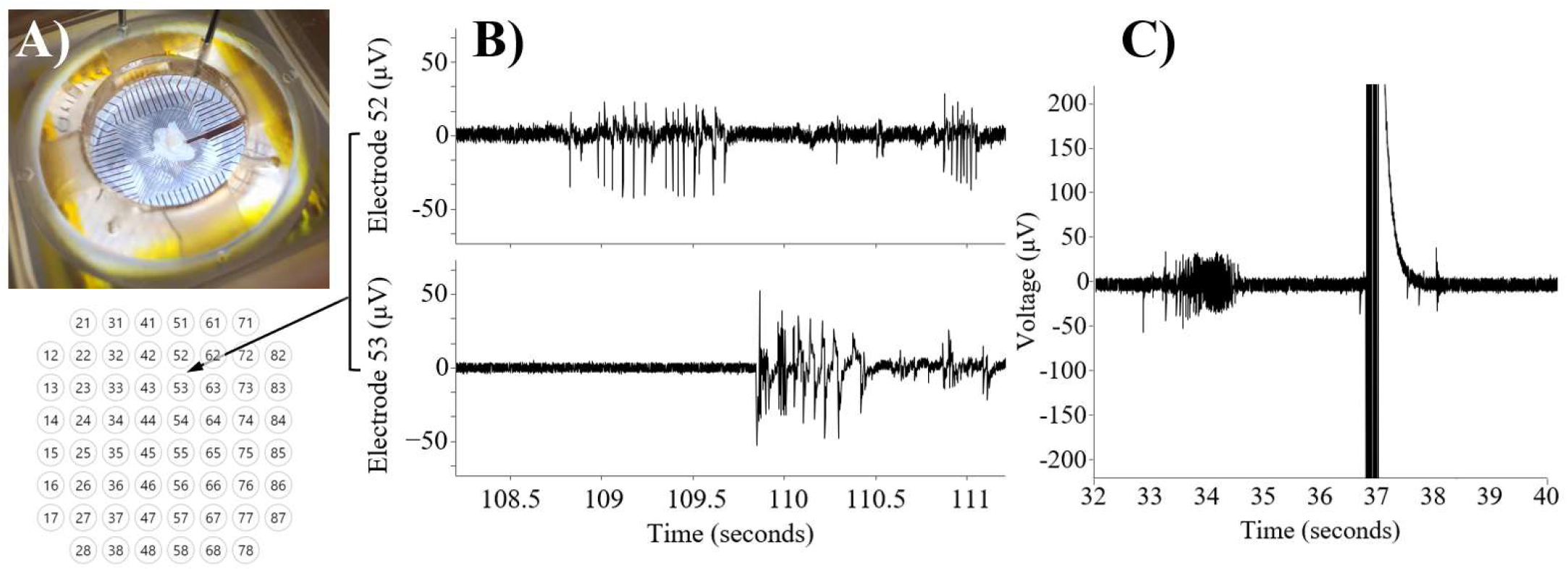
In vitro recordings. A) Rodent hippocampal slice. B) Electrophysiological recordings of pharmacologically induced seizure activity using 4-Aminopyridine (4-AP) C) Electrical stimulation applied during ictal bursting.

## X. Discussion

### A. Factors enabling OpenMEA

In recent years, key developments have laid the foundation to catalyze OpenMEA. The democratization of additive manufacturing has led to broadly accessible 3D printing using both FDM and MSLA processes which can support the mechanical and microfluidics requirements of MEA systems. The accessibility of advanced PCB manufacturing has resulted in the ability to produce the high-density features and cytocompatibility needed for MEAs. The availability of highly-capable low-noise neural interfacing ICs from companies such as Intan has allowed researchers to innovate with less reliance on custom ASICs developed in academia. Taken together, these factors have enabled OpenMEA to fulfil many of the needs of modern in vitro bioelectronics research with a cost of materials and manufacturing of <$5000 at the time of writing. This contrasts with commercial systems with a typical cost on the order of $100k.

### B. Translational device development

Collaborative platforms have been identified as the missing link in neurotechnology between benchtop development and clinical deployment [63]. OpenMEA aims to address this issue from hardware, software, and fundamental neuroscience perspectives. In closed-loop algorithm development, existing methods to solve the pathological state detection problem have ranged from thresholding of biomarkers to deep-learning approaches [64]. Analyzing the performance of these detection models is conventionally done with the use of pre-recorded (or offline) datasets. However, while offline datasets may be suitable for quantifying the efficacy of state detection algorithms, they do not consider the causal effects of how a detection-triggered stimulus applied at one point in time could change proceeding states. This is a fundamental limitation with existing approaches to closed-loop algorithm development.

Investigating the cause-and-effect dynamics of stimulation could be achieved using simulations with an online data source such as a computational model of the brain. While such models exist, they do not capture complex brain dynamics in enough detail to serve as a suitable platform for closed-loop development [65]. For this reason, quantifying the performance of a closed-loop system is must ultimately be done with biological tissue. Animal models are used as an essential translational step to validate a device performance to minimize the financial risk associated with the large investments necessary to manufacture devices and obtain regulatory approval for use with humans. To appreciate the significance of this, we must first understand the steps necessary to manufacture such medical devices and then consider how the required components could be validated with the lowest risk and overhead.

From a hardware perspective, closed-loop devices typically contain digital integrated circuits which are responsible for processing neural signals to extract biomarkers and determine an appropriate stimulus to apply in response to a detected brain state [66],[67]. Creating such ICs requires closed-loop logic to be synthesized from a hardware description language (HDL) in software by a device designer. The digital closed-loop ICs are typically connected to analog front end (AFE) ICs which are responsible for sampling the neural signals at the implanted electrode site, and delivering the electrical charge needed for neurostimulation. Although the design of AFE ICs is continuing to develop to address practical challenges (such as simultaneous recording and stimulation [68]), the key bottleneck in the efficacy of closed-loop devices is the digital control for classifying neural signals into representative brain states and generating the most appropriate neurostimulation waveform in response. For this reason, an array of digital IC architectures have been proposed [66], [69], [70]. However, few have demonstrated their efficacy in an online closed-loop setting due to the practical and regulatory challenges of *in vivo* testing. OpenMEA enables researchers to overcome this challenge and realistically quantify an IC’s efficacy with real tissue. The digital closed-loop ICs are typically connected to analog ICs which are responsible for sampling the neural signals at the implanted electrode site, and delivering the electrical charge needed for neurostimulation. To allow for the development of analog and digital ICs in isolation, the proposed solution uses modular PCBs with an FPGA prototyping device in the loop. The HDL which is used to create a low-power digital IC can be mapped to the FPGA to enable testing of closed-loop functionality *in vitro*. This approach minimizes the risks and costs of *in vivo* validation and offers an additional validation step before investing in the manufacturing of expensive application specific integrates circuits (ASICs).

### C. Towards adaptive closed-loop stimulation

Existing work on adaptive stimulation has focused on sensory adaptation (i.e. adapting to brain states). However, relatively little focus has been placed on stimulus adaptation. The bi-phasic rectangular waveform is conventionally used due to its charge-balanced property and the ease of generation with a simple current source; however, it has been shown that more complex waveforms can provide more precise temporal control of neural activity [71]. Electrical stimulators must generate waveforms while preventing damage caused by imbalanced charges during positive and negative stimulation phases. The selection of appropriate waveform parameters to evoke a desired response (e.g. seizure termination in epilepsy) is a therefore a difficult task which is infeasible to address computationally. MEAs allow for controlled *in vitro* experimental exploration of this parameter space to find the most suitable stimuli for neural control and for the treatment of disorders. This exploration is facilitated in OpenMEA through user-programmable charge-balanced stimulus waveform generators which can reside in either software of hardware.

## XI. Conclusion

The OpenMEA platform aims to ignite open-source closed-loop electrophysiology to enable breakthroughs in fundamental research and medical device development. Using accessible manufacturing processes, it enables more equitable access to bioelectronic research and development tools. This method of scientific hardware development enables a much broader audience to participate in experimentation both as research and teaching platforms when compared to existing proprietary methods.

While applications in neuroscience have been highlighted in this work, disease-on-chip models including cardiac and gastrointestinal disorders could equally leverage the OpenMEA platform. This could help to catalyze a new wave of innovation, discovery, and therapy to make electroceuticals the mainstay of medical treatment.

## XII. Author Contributions

Gerard O’Leary – Conceptualization, physical design, and assembly (FDM 3D printing and enclosure manufacturing, microfluidics MSLA 3D printing, PCB and MEA design, schematics, layout, hardware bringup), *in vitro* experiments, writing, data curation, project administration.

Iouri Khramtsov – OpenMEA studio, writing.

Rakshith Ramesh – Firmware development, writing

Prajay Shah – Microscopy.

Aidan Perez-Ignacio – Tissue preparation.

Homeira Moradi – Tissue preparation, general guidance.

Adam Gierlach – Power PCB.

Roman Genov – Conceptualization, Funding Acquisition,

Reviewing, Resources, Supervision.

Taufik Valiante – Conceptualization, Funding Acquisition,

Reviewing, Resources, Supervision.

**Supplementary Figure 9.**
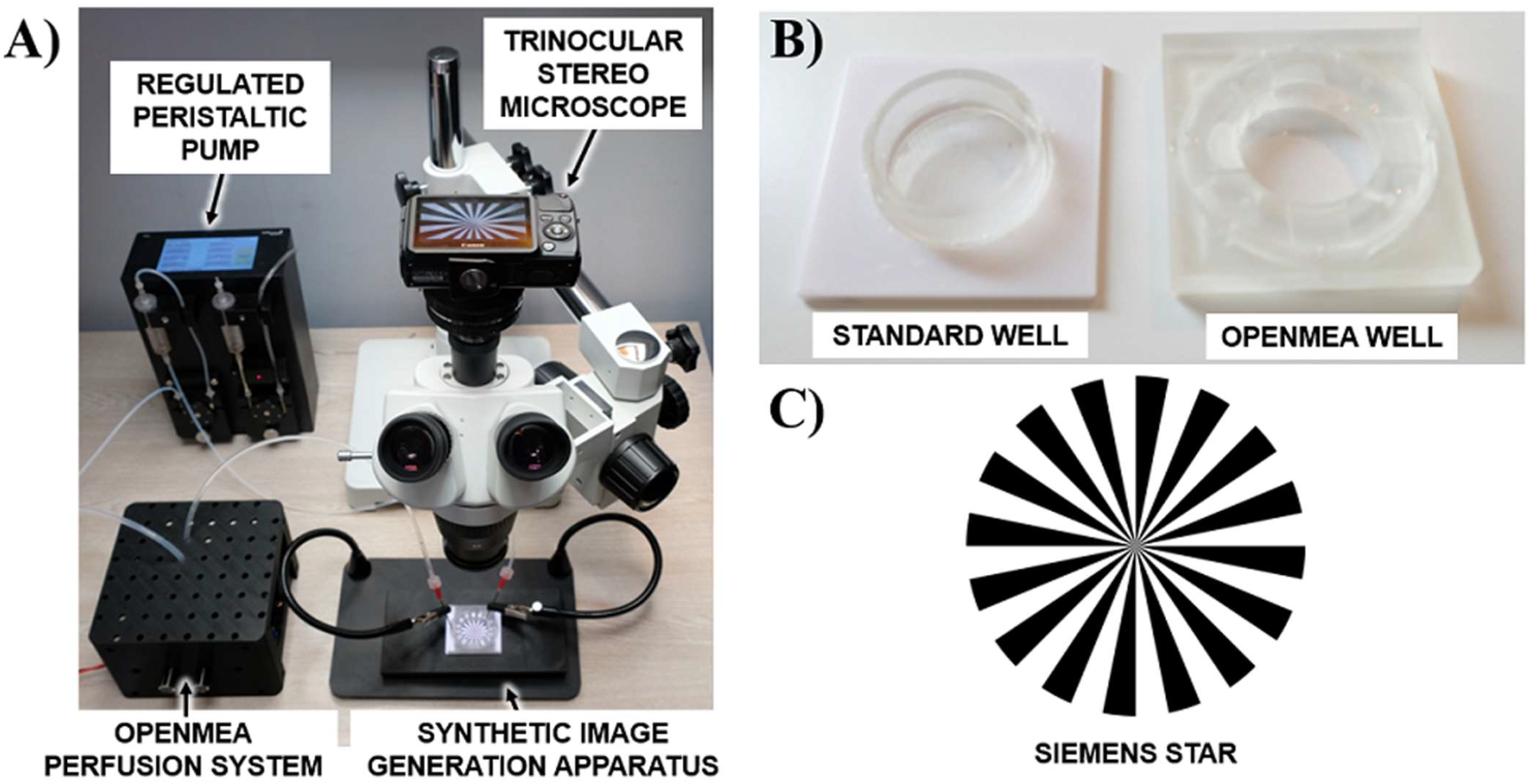
A) Focus-drift experimental test setup for analysis with synthetic test images. B) Two test MEAs with transparent bases: the first uses an 8mm high standard well, the second uses the OpenMEA microfluidics well. C) Siemens star calibration image.

## XIII. Imaging validation setup

The focus-drift prevention mechanism is validated using the test setup shown in Supplementary Figure 9. A synthetic image generation apparatus is comprised of a 3D printed MEA mounting bracket on an LCD screen with a 50µm pixel diameter (Apple iPhone 8 Plus). Two test MEAs with transparent bases are used for comparison as shown in Figure Supplementary Figure 9B; the first uses an 8mm high standard well which relies on cannulas for input perfusion and suction, the second uses the OpenMEA microfluidics approach. A standard Siemens star calibration image (Supplementary Figure 9C) is projected beneath the transparent base of each test MEA during active perfusion and video capture is performed using a trinocular stereo microscope.

**Supplementary Figure 10.**
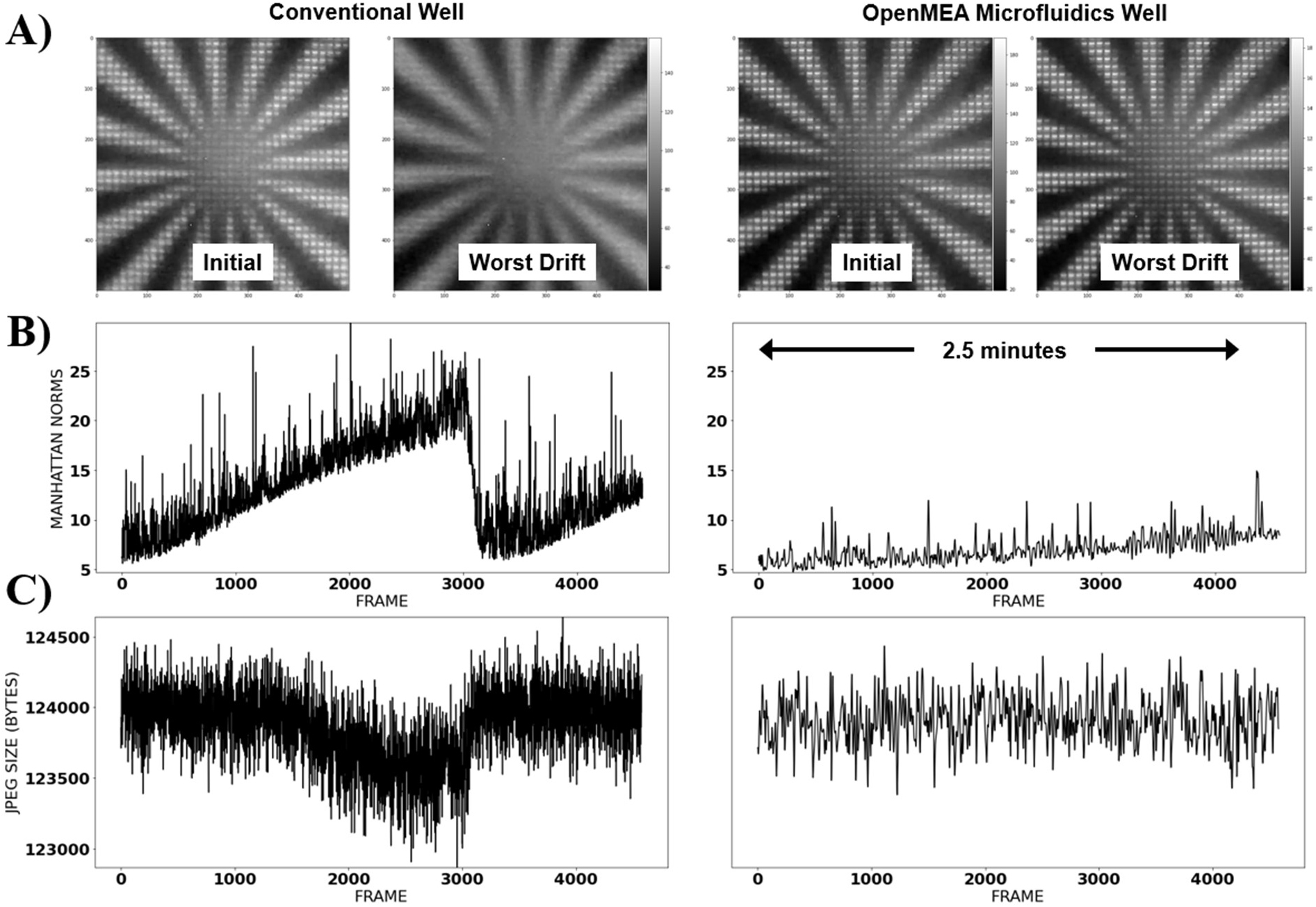
A) Initial calibrated state and worst-case focus drift for a conventional well design compared to the OpenMEA microfluidics well. B) Image drift focus quantified using the Manhattan norm. C) Image drift focus quantified using JPEG compressed data sizes.

## XIV. Imaging validation quantification

Supplementary Figure 10A illustrates the initial calibrated state, and the worst-case focus drift over a 2.5 minute period for a conventional well design compared to the OpenMEA microfluidics well. The image drift focus can be first quantified by comparing the initial calibrated image to proceeding frames in a video captured during perfusion. The Manhattan norm, *d*, is used to measure this difference as follows:

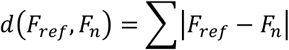

Where *F*_*ref*_ is the initial calibrated reference image, and *F*_*n*_ is the comparison frame. The difference is normalized by the number of pixels. Supplementary Figure 10B demonstrates a gradual increase in ACSF levels until the suction system overcomes the surface tension. The suction removes the ACSF to a lower level and the process repeats, causing sudden large decreases in ACSF, with a slow buildup. As the ACSF is translucent, this results in image distortion. In the case of the OpenMEA microfluidics well, these sudden transients do not occur as the ACSF is continuously removed at a slow rate via the vacuum loop. However, it is noted that there is a gradual drift which is attributed to small physical movements in the test setup. The Manhattan norm is sensitive to these physical movements which do not necessarily compromise image quality but do cause differences between the reference image and the comparison frames. A more appropriate measure of the information in each frame is the use of an entropy encoding method, which is conveniently implemented here as JPEG compression (Supplementary Figure 10). By comparing the compressed image sizes for each frame, the information content changes with the conventional well but remains stable for the OpenMEA microfluidics well.

**Supplementary Figure 11.**
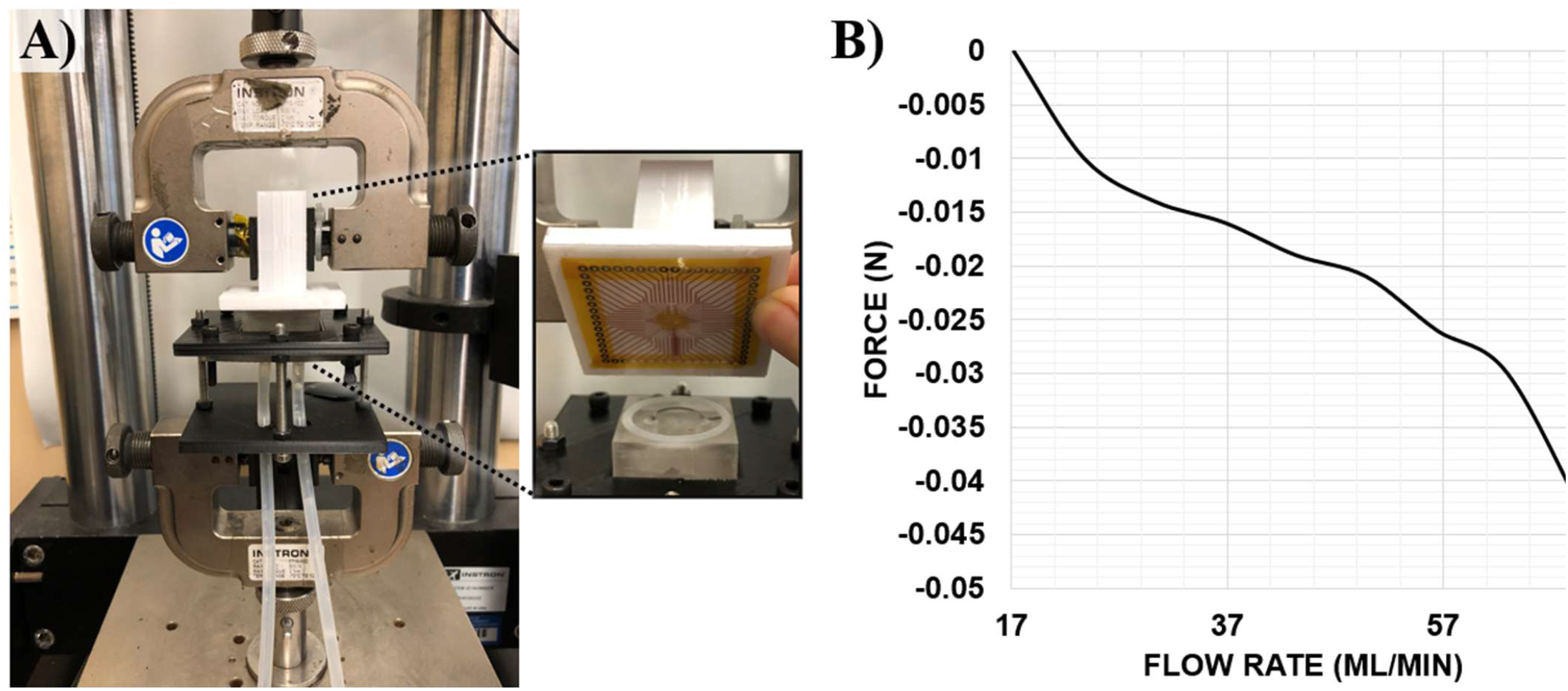
A) Bottom perfusion-based suction test setup consisting of an OpenMEA PCB adhered to a test adapter and a mechanical tensile tester. B) Increased rate of suction corresponding to an increasing flow rate in the bottom perfusion bracket.

## XV. Bottom Suction Validation

The bottom perfusion-based suction was assessed using the test setup shown in Supplementary Figure 11. An OpenMEA PCB was adhered to a test adapter to be used with a mechanical tensile tester (Instron Inc.). Supplementary Figure 11B demonstrates the increased rate of suction corresponding to an increasing flow rate in the bottom perfusion bracket, with a total suction of almost 0.05N. This is a sufficient force to ensure that tissue does not float freely within the well and is instead pulled towards the applied negative force at the electrode.

## XVI. Bottom Perfusion Validation

The ability of the suction to supply ACSF to the bottom of the well through the electrode perforations was validated using the test setup shown in Supplementary Figure 12. A green fluorescent pigment was mixed with ACSF and perfused into the well under UV light while being recorded using a trinocular stereo microscope for analysis. Supplementary Figure 12B shows the initial state of the test MEA under white light, and the state of the MEA following 60 seconds of bottom perfusion. Supplementary Figure 12C quantifies the change in green pixel values over time, showing a linear increase in pigments (and hence, ACSF) deposited in the well over the course of the experiment.

**Supplementary Figure 12.**
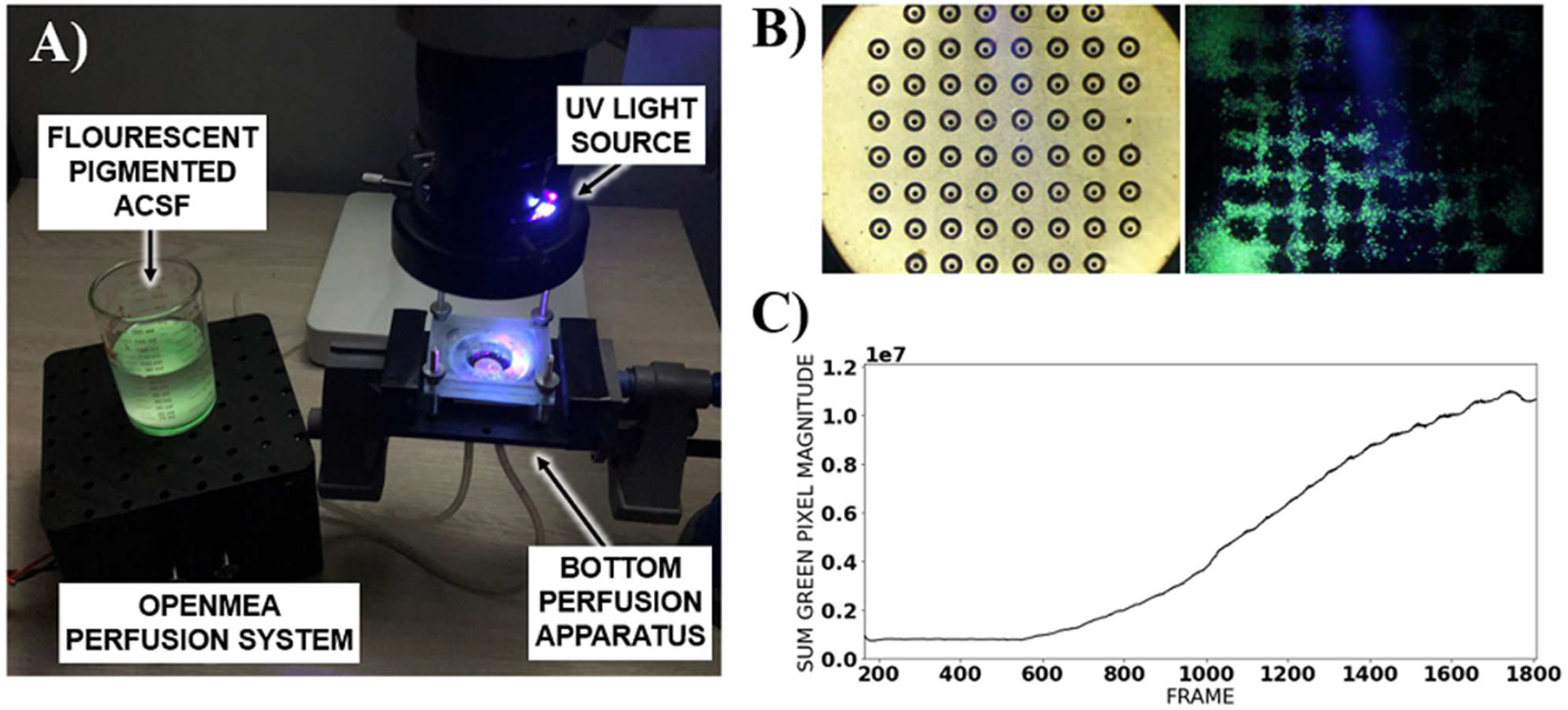
A) Bottom perfusion ACSF distribution analysis setup using green fluorescent pigment, UV light source, and trinocular stereo microscope. B) Initial state of the test MEA under white light, and the state of the MEA following bottom perfusion. C) Change in green pixel values showing a linear increase in ACSF pigments deposited in the well over time.

